# myCircos: Facilitating the Creation and Use of Circos Plots Online

**DOI:** 10.1101/052605

**Authors:** Caroline Labelle, Geneviève Boucher, Sébastien Lemieux

**Affiliations:** Institute for Research in Immunology and Cancer, Université de Montréal, Montréal, Québec, Canada; Department of Computer Science and Operations Research, Université de Montréal, Montréal, Québec, Canada.

## Abstract

Circos plots were designed to display large amounts of processed genomic information on a single graphical representation. The creation of such plots remains challenging for less technical users as the leading tool requires command-line proficiency. Here, we introduce myCircos, a web application that facilitates the generation of Circos plots by providing an intuitive user interface, adding interactive functionalities to the representation and providing persistence of previous requests. myCircos is available at: http://mycircos.iric.ca. Non registered users can explore the application through the Guest user. Source code (for local server installation) is available upon request.

## Introduction

Circos plots facilitates the visualization of sequencing data and relevant genomic information (Krzwinski et al., 2009). They are often used to integrate large amount of genomic data for one or multiple samples to emphasize particularities, similarities or differences, in a single graphical representation. However, the installation and usage of the Perl-based Circos software can be challenging for a number of researchers as it requires command-line proficiency. Other softwares have recently been developed which bypass the usage of the Perl software, among them the R packages RCircos (Zhang et al, 2013), CIRCUS (Naquin et al, 2014), circlize (Gu et al., 2014) and omicCircos (Hu et al., 2014), a Java software J-Circos (An et al., 2015), a d3-based JavaScript library BioCircos.js (Cui et al., 2016) and a web-based interactive service ClicO FS (Cheong et al., 2015). Nevertheless, most of these alternatives still require coding efforts from the user or expect some knowledge in applied bioinformatics.

Here, we describe a new web application that enables the creation of Circos plots in a user-friendly manner: only formatted data files are required from the user. We opted for a simplified interface resulting in a default visualization that can be easily modified later on. Our web-based interface also provides useful functionalities such as: a database that stores previously generated plots, and interactive SVG (Scalable Vector Graphics) output. Extending the typical and static graphical output, users can interrogate their plots using the interface, directly within their web-browser. We decided to integrate the Perl-based Circos tool in our application to maximize the compatibility between myCircos and the original tool (using our generated configuration files in a console environment or using existing configurations in our application).

## Application

### Usage

myCircos is publicly accessible and offers guest and registered users basic functionalities. Two options are available to create a Circos plot. In the first one, *From data file(s)*, a simplified form is filled by the user to customize the different tracks and upload its data. The data input files should be tab-delimited (.txt files) and contain the interesting genomic data (e.g. a value or a label) associated to given chromosomal locations. The organism, the number and types of tracks are among the parameters set in the form. This option can serve as a first initial step. The second option, *From configuration files*, is useful when a user already have Circos configuration files (by going once through the form or by other means) and wants to take advantage of the added interactivity offered by our application. The user provides a package (e.g. ˙zip) containing the needed Circos configuration and tab-delimited data files.

It typically takes several minutes to generate the image depending on the amount of data submitted and the number of tracks required. Registered users benefit from myCircos’ database functionalities: their webbrowser can be closed as the output will be automatically and safely stored. It will be accessible when ready via the “myCircos” menu item and the user will be notified by e-mail. In addition, registered users’ have access their previously created plots and can revisit or delete them.

### Advantages and Features: Database and Interactivity

The registered user’s list of plots is found under the “myCircos” tab. For each plot, basic information (e.g. miniature thumbnail, creation date, experiment description) are provided. The plot (as an SVG file) can be downloaded from this page as well as the configuration package when available (i.e. for plots generated with the “From data files” option). By clicking on a given Circos plot, the SVG output is loaded in the web-browser, allowing interactions. Such interactivity makes it easier to further analyze the data. By zooming and moving, a user can focus on a particular region of the plot. By clicking on a feature with the “Highlights” option enabled, a user can emphasize elements of interest. Lastly, with the “Value & Gene Information” option enabled, it is possible to access the associated data point value by hovering on a graph feature, and to access information about the genes overlapping the selected feature by clicking on it (Figure1 A). If a single gene overlaps a feature of interest, useful information about this gene is displayed in the gene information box. When multiple genes are overlapping, the number of overlapping genes is presented. A legend and a detailed information section also appear on the page for each graph as well as links to download the package and SVG files.

**Figure 1.**
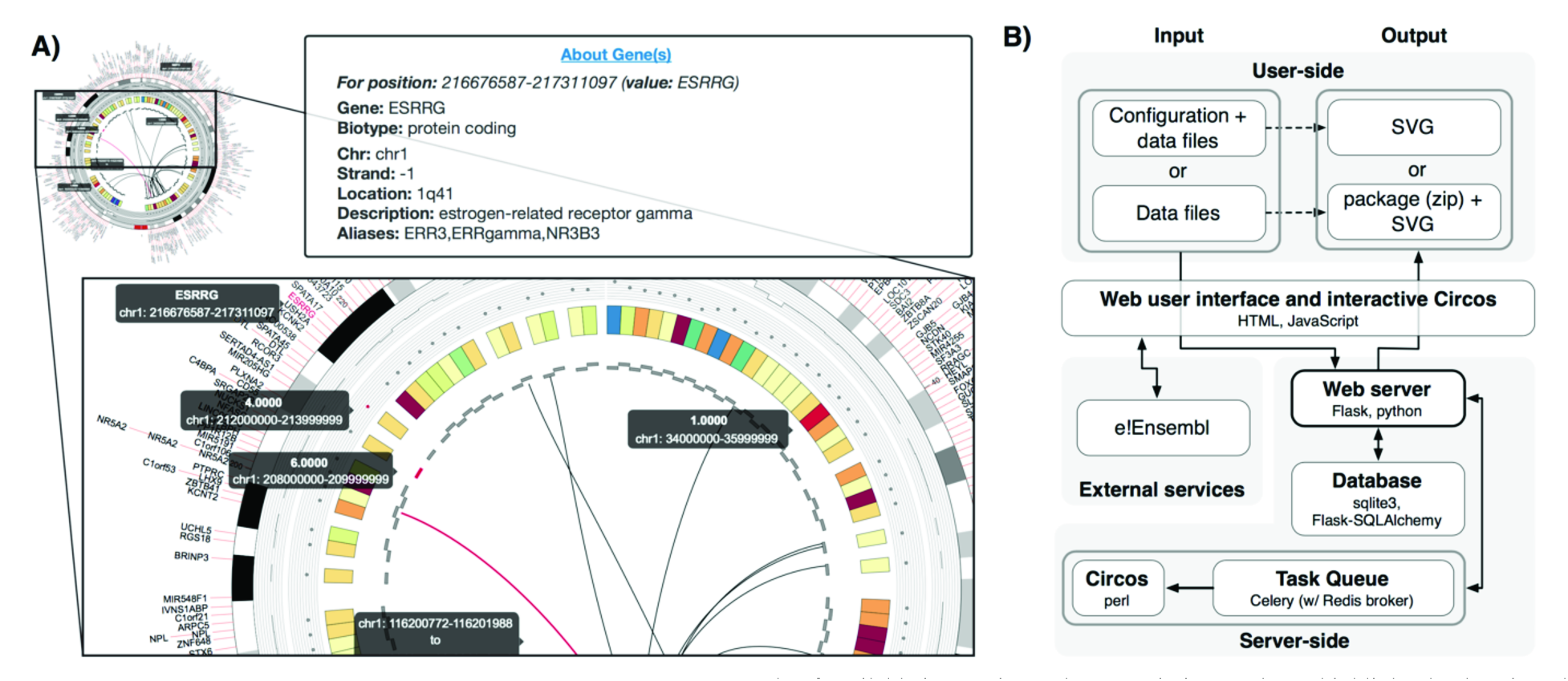
myCircos’s interactivity and client-server architecture. **A**. Example of available interaction. The zoom-in image shows highlighted values in red on the various tracks. Value-information tooltips display data input values for the selected elements as well as gene information for their related chromosomal position (gene information box;upper-right). **B**. Overview of the client-and server-side components. The web interface feeds the user input to the web server, which calls the Perl-based Circos software within a Celery task queue before returning the user-output file

## Implementation

Figure 1B describes the client-server architecture implemented using, on the server side: Circos (Perl-based command-line tool), Flask (Python-based micro framework, http://flask.pocoo.org/), Flask-SQLAlchemy extension (object-relational mapper, http://flask-sqlalchemy.pocoo.org/), SQLite (registered user’s profiles, https://www.sqlite.org/), Celery (distributed task queue system, http://www.celeryproject.org/), Redis (as Celery’s broker, http://redis.io/). Gene information are obtained by querying the e!Ensembl RESTful API (http://rest.ensembl.org/) and the NCBI Entrez Utilities API (http://www.ncbi.nlm.nih.gov/books/NBK25501/). Provided data, such as the karyotype for human and mouse, is derived from UCSC Genome Browser data tables. On the client side, standard HTML5 and JavaScript is used as well as Jinja (templating engine, http://jinja.pocoo.org/) and Bootstrap (http://getbootstrap.com/). Interactivity is mainly provided by the use of jQuery (https://jquery.com/).

## Conclusion

myCircos greatly facilitates the creation of Circos plots and complements the original tool with interactive visualization. Being flexible and user-friendly but still exposing the key features of original command-line tool, it removes some of the challenges associated with Circos plot creation.

### Funding

This work has been supported by the Institute for Research in Immunology and Cancer [IRIC Next Generation Award to C.L.].

### Conflict of Interest

none declared.

## Acknowledgements

The authors would like thank Vincent-Philippe Lavallée, Alexandre Rouette, Diana Paola Granados and Patrick Gendron for their comments and suggestions.

